# Measuring factors affecting honey bee attraction to soybeans using bioacoustics monitoring

**DOI:** 10.1101/2022.11.04.512777

**Authors:** Karlan C. Forrester, Chia-Hua Lin, Reed M. Johnson

## Abstract

Soybean is an important agricultural crop around the world, and previous studies suggest that honey bees can be a component for optimizing soybean production through pollination. Determining when bees are present in soybean fields is critical for assessing pollination activity and identifying periods when bees are absent so that bee-toxic pesticides may be applied. There are currently several methods for detecting pollinator activity, but these existing methods have substantial limitations, including the bias of pan trappings against large bees and the limited duration of observation possible using manual techniques. This study aimed to develop a new method for detecting honey bees in soybean fields using bioacoustics monitoring. Microphones were placed in soybean fields to record the audible wingbeats of foraging bees. Foraging activity was identified using the wingbeat frequency of honey bees (234±13.9 Hz) through a combination of algorithmic and manual approaches. A total of 243 honey bees were detected over ten days of recording in four soybean fields. Bee activity was significantly greater in blooming fields than in non-blooming fields. Temperature had no significant effect on bee activity, but bee activity differed significantly between soybean varieties, suggesting that soybean attractiveness to honey bees is heavily dependent on varietal characteristics. Refinement of bioacoustics methods, particularly through incorporation of machine learning, could provide a practical tool for measuring activity of honey bees and other flying insects in soybeans as well as other crops and ecosystems.

## Introduction

### Honey Bees and Soybean Production

Soybean (*Glycine max* (L.) Merr.) is a growing agricultural crop around the world and has become the number one agricultural export of the United States. In 2020, 17.6% of agricultural export value in the United States was attributed to soybeans (USDA FAS 2021). The United States exported $25.7 billion worth of soybeans grown for oil, animal feed, industrial products, and human consumption (USDA FAS 2021), an increase of $7.0 billion from the previous year (USDA FAS 2020). In 2021, soybeans covered 87.6 million acres of agricultural land in the United States, an increase of 5% from the previous year (NASS, USDA 2021). Soybeans are also a major crop in Ohio. Of the 9.1 million acres of principal crops planted in Ohio during 2021, 4.8 million acres (52%) were soybeans. This accounted for 5% of the nation’s annual soybean production (NASS, USDA 2021).

The growing soybean industry is faced with the continued need to optimize resource input and maximize yield output, and many studies suggest that honey bees (*Apis mellifera* Linn.) can be a component for optimizing soybean production through pollination. Soybeans are self-fertile angiosperms that often exhibit cleistogamy, meaning that self-fertilization occurs within closed flowers (Benitez et al 2010). Because of this, early publications purported that soybeans are not attractive pollinator plants and are rarely visited by honey bees (Lent 1934, Milum 1940). However, more recent studies have shown that honey bees frequently forage in soybean fields, and cross-pollination by pollinators contributes additional benefits to soybean fruiting. Soybean yield is positively correlated with honey bee visitation, with yield increases ranging from 5.7% to 81% across studies, likely due to differences in experimental methods, soybean varieties, and variables such as time and location (Blettler et al 2018, Chiari et al 2005a, Erickson 1975a, Erickson et al 1978, Esquivel et al 2021, Issa et al 1984, Jaycox 1970, Juliano 1976, Kettle & Taylor 1979, Levenson et al 2022, Milfont et al 2013, Monasterolo et al 2015, Santos et al 1993, Toledo et al 2011, Vila et al 1992). Chiari et al (2005a) demonstrated that pollination by honey bees and pollination by wild pollinators both increased soybean yield. Milfont et al (2013) demonstrated a yield increase of 11% when open-pollinated soybeans were supplemented with nearby honey bee colonies. Soybean yield also may benefit from the presence of nearby pollinator habitat (Levenson et al 2022). Together, these studies suggest that optimal soybean production can be achieved through the combined pollination activity of wild pollinators and managed honey bees. Milfont et al (2013) estimated that honey bee pollination of soybeans could be contributing $59.70 to $110.50 of profits per hectare, representing a contribution of $6.1-17.4 billion to the global economy.

Honey bee pollination can have multiple effects on soybean fruiting that may lead to increased yields. Pollination has been shown to increase hybridization (Abrams et al 1978, Chiang & Kiang 1987, Cutler 1934, Jaycox 1970, Sim & Choi 1993), number of pods (Chiari et al 2005a, Cunningham-Minnick et al 2019, Erickson et al 1978, Juliano 1976, Moreti et al 1998, Pinzauti & Frediani 1980), number of seeds (Blettler et al 2018, Cunningham-Minnick et al 2019, Moreti et al 1998, Pinzauti & Frediani 1980), pod weight (Juliano 1976, Toledo et al 2011), fruiting rate (BeaudelaineKengni et al 2015, Sim & Choi 1993, Tchuenguem & Dounia 2014), healthy seed (BeaudelaineKengni et al 2015, Tchuenguem & Dounia 2014), and seeds per pod (BeaudelaineKengni et al 2015, Issa et al 1984, Milfront et al 2013, Moreti et al 1998, Pinzauti & Frediani 1980, Tchuenguem & Dounia 2014). Pollination has also been shown to decrease seed abortion (Erickson et al 1978, Monasterolo et al 2015) and flower abortion (Chiari et al 2005b, Monasterolo et al 2015).

Some of the earliest studies on the relationship between honey bee pollination and soybean yield reported that yield benefits differ by variety, likely due to variation in bloom characteristics and their resulting attractiveness to honey bees (Erickson 1975a, Issa et al 1984). Differences between soybean varieties include flower color (ranging from white to violet), growth habit (determinate or indeterminate), maturity group (time from planting to maturity), frequency of cleistogamy, flower size, flower fragrance, number of flowers, nectar volume, and nectar sugar concentration (Benitez et al 2010, Erickson 1975b, Stowe & Vann 2022). In addition to phenotypic variation across soybean varieties, soybeans produce nectar of varying quality and quantity depending on environmental factors, including temperature (Robacker et al 1983), time of day (Blettler et al 2016, Severson & Erickson 1984), soil macronutrients (Robacker et al 1983), and soil microbiota (BeaudelaineKengni et al 2015).

Variations in soybean bloom characteristics are correlated with attractiveness to honey bees. Sugar concentrations above 25% and low rates of cleistogamy are generally most attractive to honey bees (Erickson 1975b), whereas flower color has little impact on honey bee foraging (Chiang & Kiang 1987, Jaycox 1970, Mason 1979, Severson & Erickson 1984). Attractiveness also changes over time, with most studies observing peak honey bee activity in soybeans around midday (BeaudelaineKengni et al 2015, Blettler et al 2016, Chiari et al 2005a, Issa et al 1984, Jaycox 1970, Santos et al 1993, Toledo et al 2011). This is likely due to changes in floral nectar quality and quantity throughout the day, with sugar concentration increasing and nectar volume decreasing as the day progresses (Severson & Erickson 1984). Honey bee foraging also peaks in the middle of the month-long blooming period when flower production is at its maximum (Blettler et al 2016).

### Detecting Honey Bee Activity

Because soybeans are capable of self-pollination – and possibly as a consequence of early studies purporting that honey bees are poor pollinators of soybeans – honey bees are often disregarded as a factor of production, which opens the door for potentially bee-toxic insecticides to be applied during soybean bloom (Jaycox 1970, Milfont et al 2013). Many soybean insecticides applied during bloom are highly toxic to bees and carry cautionary language to protect bees on the pesticide label. While there have been reports that pyrethroid insecticides repel bees from areas where they have been applied (Rieth & Levin 1988), the pyrethroids cypermethrin and lambda-cyhalothrin were not found to deter honey bee foraging when applied to blooming soybeans (Fagúndez et al 2016). Pesticides are picked up by foraging honey bees and cause lethal and sub-lethal effects in the colony, creating a major risk for foraging colonies and reducing long-term viability of pollination and its associated benefits (Reichelderfer & Caron 1979, Santos et al 2013). This both raises legal concerns and prevents growers from fully implementing honey bee pollination into integrated pest and pollinator management (IPPM) frameworks. The conditions for pesticide application in soybeans must be reassessed within the context of honey bee activity to support the long-term sustainability of soybean production.

In order to assess how environmental variables and soybean varieties can predict honey bee foraging in soybeans, we must first be able to detect foraging honey bees across time. There are several existing methods for detecting pollinator activity, mainly through the use of pan traps, visual observation, and manual collection (Portman et al 2020). Pan traps are brightly colored bowls filled with soapy water that are used as a passive sampling method. Pollinators are attracted to the bright colors, then become trapped in the liquid. Manual collection methods include targeted netting and sweep netting, and visual observation provides a nonlethal method of insect detection. Pan traps, visual observation, and netting have all previously been used to assess honey bee activity in soybean fields, with visual observation being the most common method (Supplemental Table 1).

Existing methods for bee detection have drawbacks that make them ineffective for accurate large-scale detection of honey bees in crops. Pan traps tend to favor smaller bees, such as those belonging to the family *Halictidae*, over larger bees such as honey bees (Portman et al 2020). Visual observation and netting techniques are time and labor intensive, and these methods are more susceptible to collector bias (Portman et al 2020, Westphal et al 2008). In addition, detection methods such as trapping are fatal to the target organisms. A novel method is necessary for accurate bee detection, particularly in large cropping systems like soybean.

Bioacoustics is a branch of science that focuses on sound production by living organisms, and bioacoustics monitoring is a relatively new method for detection and identification of species through audio recording and analysis. Deep learning approaches to bioacoustics monitoring are currently being developed for detection and identification of mosquitoes that vector malaria (Hassall et al 2021, Khalighifar et al 2022, Kim et al 2021, Kiskin et al 2020a, Kiskin et al 2020b, Vasconcelos et al 2019, Vasconcelos et al 2020). Bioacoustics monitoring has also been combined with machine learning to automate analysis of birdcall recordings and increase bird monitoring efficiency through the use of the deep neural network BirdNET (Kahl et al 2021, Toenies & Rich 2021, Wood et al 2021). A study by Zhang et al (2017) proposed the use of wingbeat spectrum imaging for insect identification using convolutional neural networks. Similar methods could be used for in-field detection and identification of any insect with a distinct and detectable wingbeat, and honey bees fulfill these criteria by producing an audible wingbeat frequency of 234±13.9 Hz (Clark et al 2017).

The goal of this study was to develop an effective and efficient bioacoustics method for detecting honey bee activity in soybean fields. To accomplish this, we made audio recordings in soybean fields around the blooming period and used a combination of automated and manual techniques to identify honey bee activity.

## Methods

### Study Area

This study was completed in four soybean fields near Apple Creek, Ohio (Supplemental Figure 1). Fields A and B were planted with the soybean variety Synergy 9720, field C was planted with Synergy 9727, and field D was planted with Synergy 9723 (Supplemental Table 2). Data was collected over ten days on July 21-24, July 26-28, and August 5-7, 2021, except for field B which yielded no data on August 7 due to a microphone malfunction. For this study, a field was considered to be in bloom during growth stages R2 and R3, and audio was recorded during stage R3. Fields A, B, and D were in bloom July 21-24 and July 26-28, while only field C was in bloom August 5-7. All four fields were located in predominantly agricultural areas and were surrounded by corn fields, wheat fields, alfalfa fields, additional soybean fields, deciduous forestland, and major and minor roads. An apiary with approximately ten colonies was located 25 meters east of field B, but the locations of other managed apiaries or feral colonies established in wooded areas were not known.

### Data Collection

Audio recorders (Sony model ICD-PX370) were attached to 0.635 × 91.44 cm (0.25 × 36 inch) square wooden stakes and protected from wind and rain with high-density foam and 3D-printed white plastic rain covers (https://www.thingiverse.com/thing:5380356, Figure 1). The recorders were programmed to record audio at the highest microphone sensitivity setting and a bitrate of 48 kbps (mono). A recorder was placed in each field 35 meters from an easily accessible field edge. The recorders were left to continuously record environmental audio for three-day periods, then retrieved for battery replacement. The location of each recorder was marked with a colored flag on top of a stake to aid in retrieval. In blooming fields, the recorders were placed at the same height as the highest blooming nodes beneath the leaf canopy. If the field was not in bloom, the recorders were placed at the same height as the highest non-blooming nodes beneath the leaf canopy. Each recorders’ radius of detection for bee wingbeats was approximately one meter.

**Figure 1.**
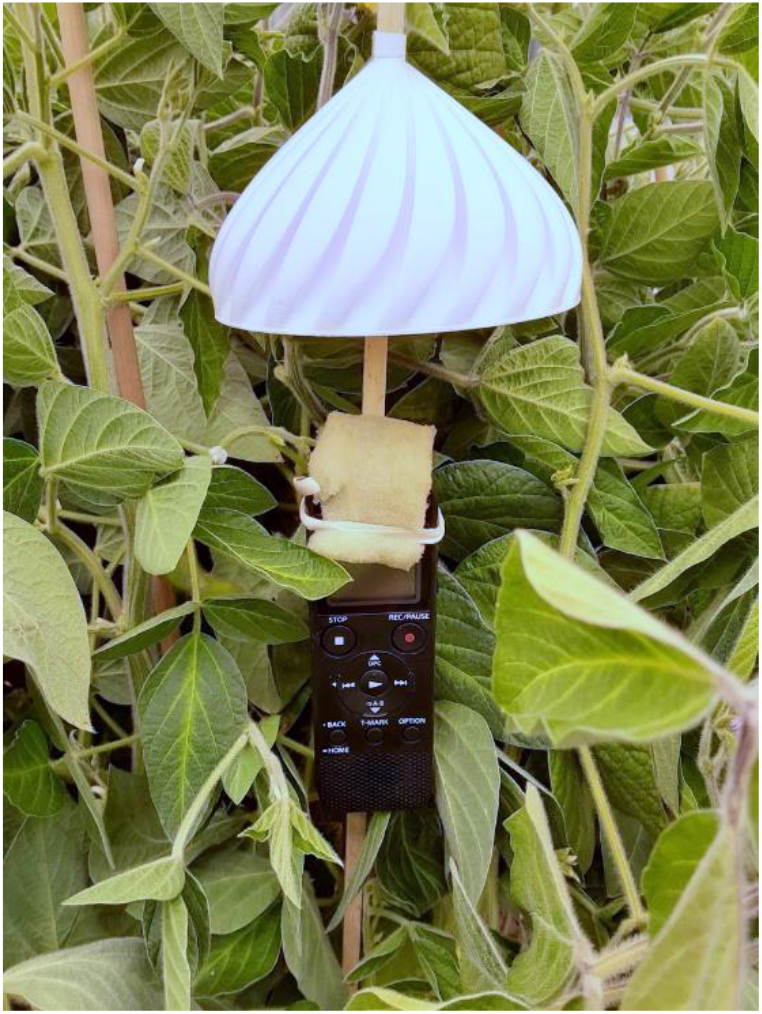
Audio recorder in soybeans.

### Audio Processing

Audio files were downloaded from audio recorders in MP3 format, downsampled to 16kHz and converted to WAV files using FFMPEG (Tomar 2006), then split into one-hour segments using the AudioSegment module in the pydub library (v.0.25.1, Robert 2018) using Python (v.3.9.7). Hours of audio recorded before sunrise or after sunset were removed from analysis. Potential bee detections were identified based on a range of audio frequencies corresponding with the second harmonic of the honey bee wingbeat frequency (370-570 Hz) with a duration of 1 sec. and a threshold of 0.0001 using the “find_rois_cwt” function in the scikit-maad soundscape analysis package in Python (Ulloa et al 2021). Possible bee detections were output as a CSV file for manual audio assessment.

### Manual Audio Assessment

Automated honey bee detections were loaded as labels overlayed on a spectrogram of one hour audio sequences and the source of each detected sound was manually identified using Audacity® (v.3.1.3, Audacity Team 2022). Audio assessment was completed by listening to the areas of interest and identifying the source of each detection event in a manually curated label file (Supplemental Table 3). Visual assessment of spectrograms in the areas of interest augmented audio assessment (Figure 2). The spectrograms were adjusted to show recorded frequencies in the 100-600 Hz range, which included the target frequency for honey bee wingbeats (234±13.9 Hz) and its second harmonic (468±27.8 Hz). Gain was set to 35 dB and spectrograms were displayed using Mel scaling to more clearly distinguish target frequencies from background noise. A total of 403 hours of audio recordings were manually assessed.

**Figure 2.**
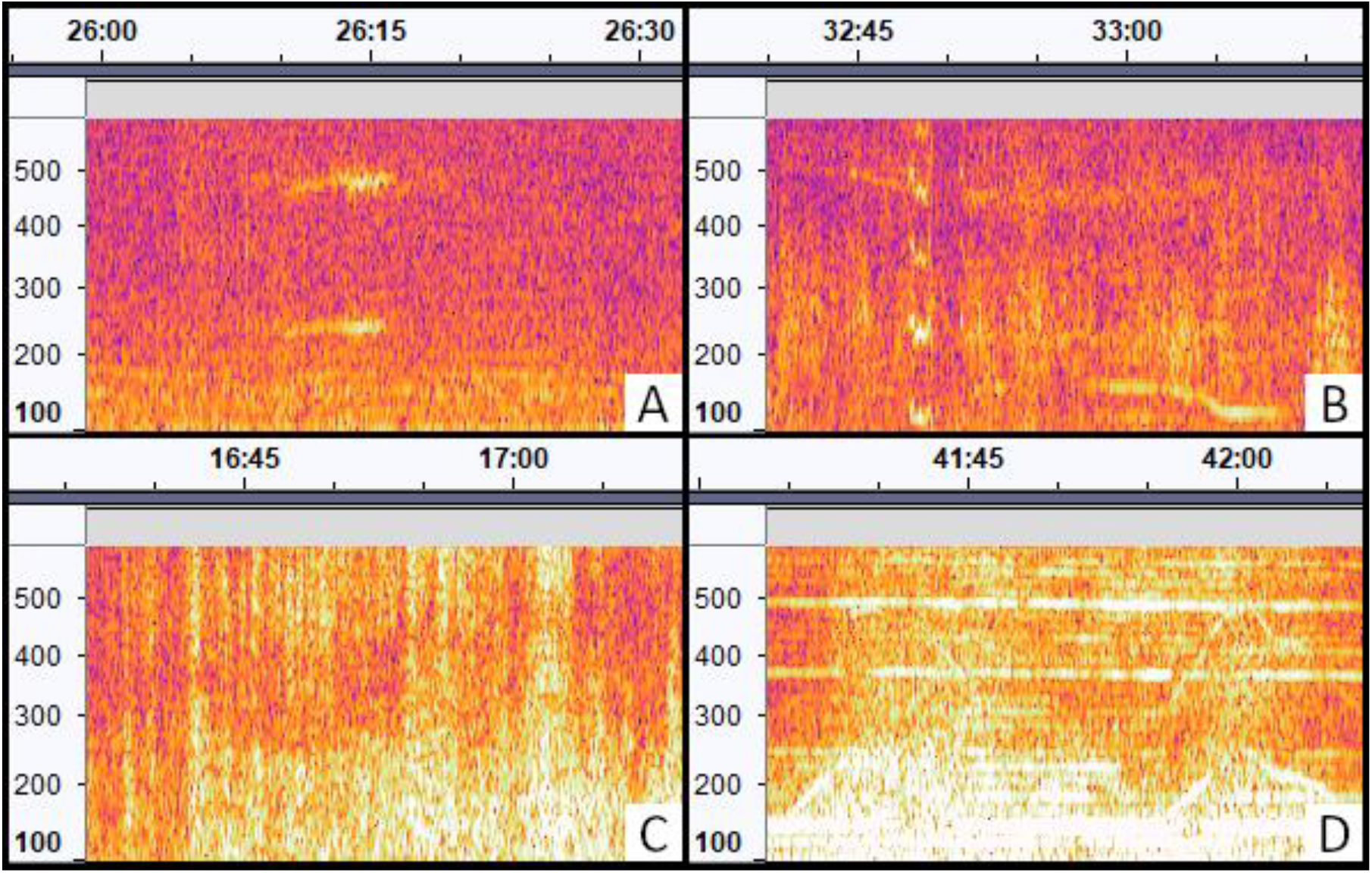
Spectrograms of potential bee activity. The x-axis shows time (mm:ss) and the y-axis shows frequency (Hz). Screenshots modified from Audacity® (Audacity Team 2022). **A)** Honey bee wingbeats (230-250 Hz) with second harmonic (460-500 Hz). **B)** Non-honey bee insect wingbeats (105-125 Hz) with second, third, and fourth harmonics. **C)** Air traffic. **D)** Agricultural equipment operating in a neighboring field.

### Statistical Analysis

Due to non-normal distribution, a Kruskal-Wallis one-way analysis of variance and Dunn’s test were used to determine significant differences in the number of bee detections between soybean varieties, between fields, and between blooming and non-blooming soybeans. Weather data for the ten recording dates were obtained from the Ohio State University CFAES Weather System Wooster Station (OSU CFAES 2022), and the correlation between average daily temperature and the number of bee detections was determined using a linear regression analysis and Pearson’s correlation coefficient. All statistical analyses were completed using R statistical software (v.4.0.3, R Core Team 2022) and visualized with the ggplot2 package (Wickham 2016).

## Results

The automated audio assessment identified a total of 11,638 potential bee detections over the ten days of recording (Supplemental Table 4). Of those detections, 10,307 were produced by vehicles, 130 were produced by insects other than honey bees, and 243 were produced by honey bees. Most bee activity occurred between 10 AM and 5 PM Eastern Daylight Time, with the greatest activity occurring between 1 PM and 4 PM (Figure 3). Field A had the least amount of recorded bee activity with only 34 bee detections. Fields B and C yielded 52 and 45 bee detections, respectively. Field D yielded 112 bee detections, more than twice the bee activity of any other field. July 21 was the most active day for bees in all three fields blooming at that time (Figure 4). Bee activity was significantly greater in blooming fields than in non-blooming fields (p = 0.001, N = 39, df = 1). Bee activity also differed significantly between soybean varieties (p = 0.004, N = 39, df = 2), with less activity in variety 9720 than in varieties 9723 and 9727, and between fields (p = 0.010, N = 39, df = 3), with less bee activity in fields A and B than in fields C and D. There was no significant correlation between bee detections and average daily temperature (p = 0.107, N = 24, df = 22).

**Figure 3.**
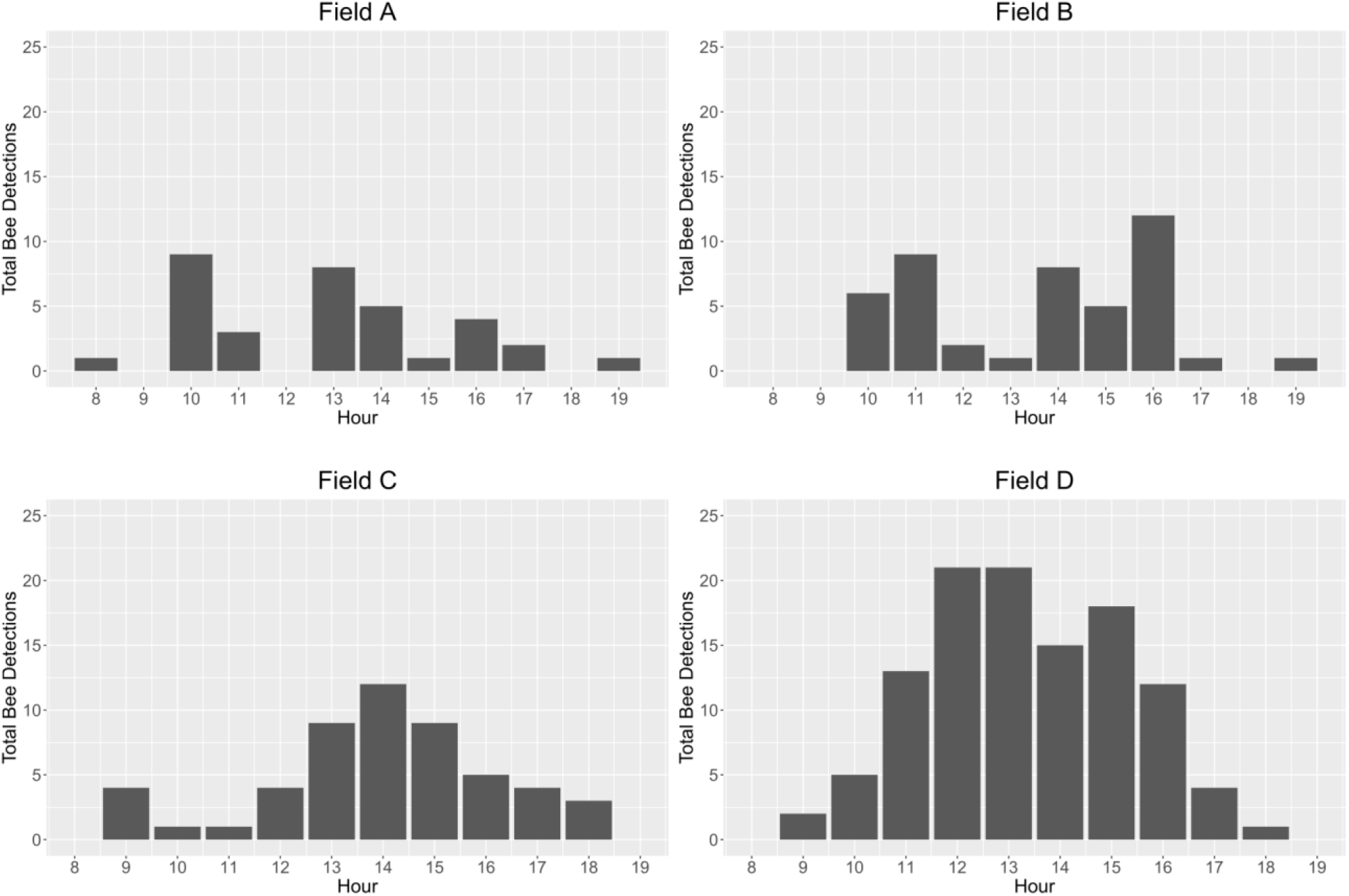
Bee detections per hour across the four study fields pooled over the 10 days of recording (July 21-24, July 26-28, and August 5-7, 2021).

**Figure 4.**
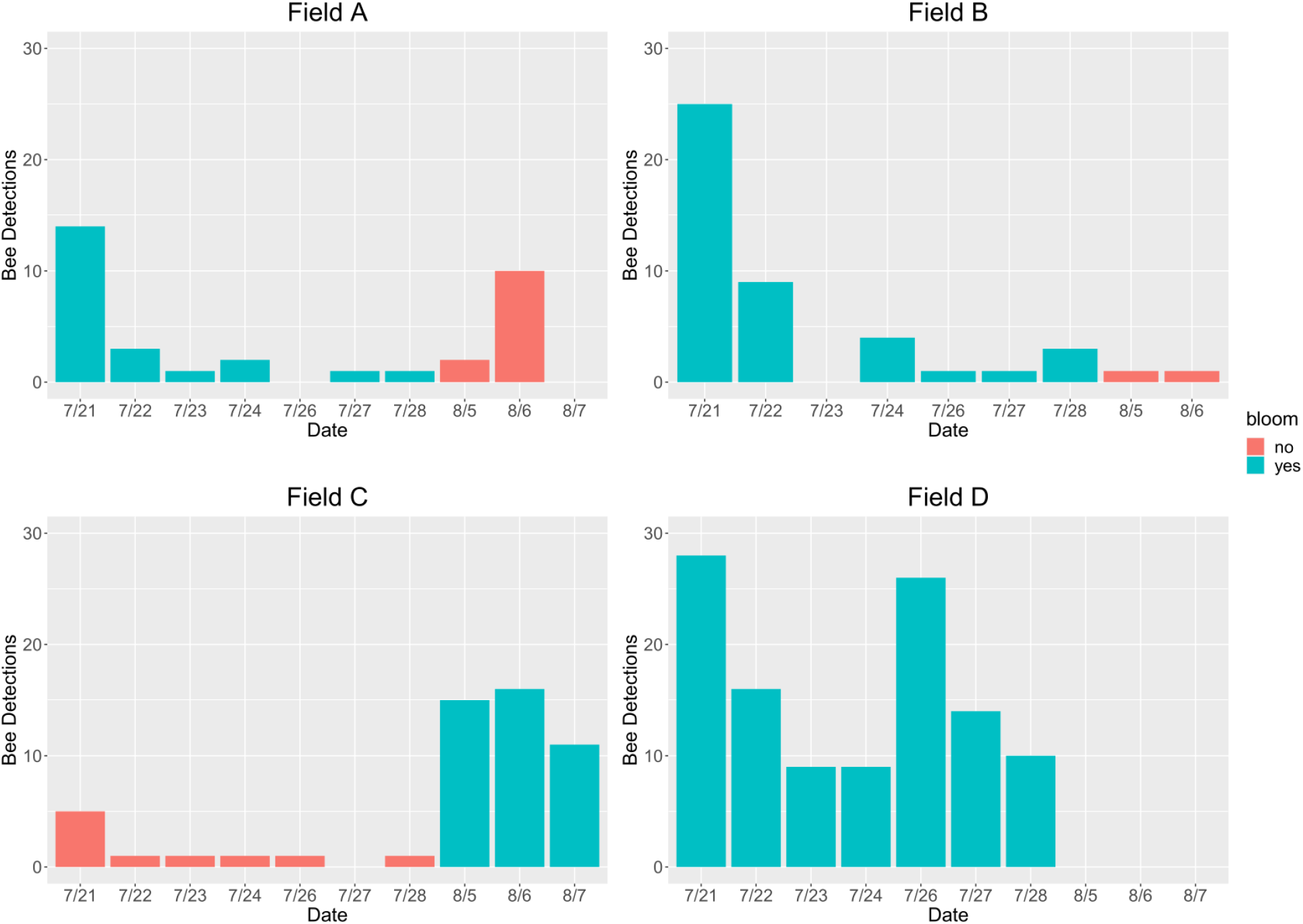
Daily bee detections in each field across all recording dates. No recording was made in field B on August 7.

## Discussion

These results demonstrate that bioacoustics monitoring is a viable method for detecting honey bees and other insects in soybean fields. Comparison of detections during bloom and non-bloom showed that there was more bee activity in each field when it was blooming than when it was not blooming. There were significant differences in bee detections between soybean varieties, with less bee activity in both fields that contained variety 9720.

There was no significant correlation between bee detections and average daily temperature. Although a study by Robacker et al (1983) found that nectar production in soybeans increased with temperature, a different study by Severson and Erickson (1984) found that temperature had no significant effect on nectar production. It is possible that temperature did not impact nectar production in the soybean varieties used in this study and thus had no significant effect on honey bee activity.

It was noted that bee detections in field A exceeded the expected bee activity for non-blooming fields on August 6, with a total of 10 bee detections. Upon reassessment of the data, it was discovered that 7 of those 10 bee detections occurred in the span of 25 seconds and were likely produced by the same bee. If this is the case, only four bees were detected in field A on this date, which more closely aligns with the number of bee detections on other dates and in other fields during non-bloom. This was the only instance of an individual bee exceeding 3 detections, and most bees were detected only once.

The recorders picked up a wide array of non-bee activity such as traffic, birds, cicadas, and agricultural equipment. By targeting the second harmonic of honey bee wingbeats, the scikit-maad soundscape analysis package was able to exclude cicadas, most birds, and other general noises from detection output. Vehicular noise occupied a similar frequency range to honey bee wingbeats, but the spectrograms produced for vehicular audio were visually distinct from the spectrograms produced for honey bee wingbeats. Despite the high occurrence of traffic detections and the proximity of some fields to major and minor roads, bee activity was still detectable, with only 7 of the 243 manually assessed bee detections overlapping with traffic detections. The only noise source that confounded bee detection was the use of agricultural equipment near fields A and C during periods of July 21, field C during periods of July 22, fields C and D during periods of July 27, fields B and D during periods of July 28, and fields C and D during periods of August 5. Bees could not be detected when agricultural equipment was present due to the sustained duration and intensity of the noise overlapping with bee wingbeat frequencies. However, agricultural equipment was only present for one or two hours on a given day, and enough bees were detected on days when agricultural equipment was present to yield usable data.

This method also proved to be much more time and labor efficient than other methods. It took approximately 8 hours to set up and retrieve the microphones across all recording periods and 34 hours to manually assess the audio data. Altogether, a total of 42 hours was needed to collect and analyze 403 hours of audio recordings. This method also allowed data to be collected in four fields simultaneously, was nonlethal, and was not susceptible to observer bias. Refinement of this method using machine learning for bee detection could further reduce the amount of time and labor required for manual audio analysis.

These results provide a general picture of when honey bees are most active in soybean fields, and conclusions can be made about when to make pesticide applications. Bee activity greatly decreases after 5 PM in blooming fields, and bees are rarely present in non-blooming fields. Based on these data, pesticide applicators can minimize honey bee exposure to harmful insecticides by only spraying soybean fields when bees are not actively foraging a) before and after the blooming period or b) after 5 PM during the blooming period.

A novel methodology such as audio detection could be used to develop a better IPPM framework for soybean growers that takes honey bee activity into account. Tracking honey bee activity in soybeans across the hours of the day has already been proposed as a means for reducing honey bee exposure to harmful pesticides (Blettler et al 2016), but this has not been utilized at a commercial scale due to the practical challenges presented by current honey bee detection methods. Bioacoustics monitoring could provide an efficient, effective, and accessible method for determining when honey bees are present in blooming soybean fields, allowing pesticide applicators to better follow pesticide label guidelines and mitigate pesticide exposure. Research to refine this audio detection technology for future implementation in precision agriculture, including the use of advanced machine learning to assist with data analysis and interpretation, is ongoing.

## Acknowledgements

This work was supported by the Agriculture and Food Research Initiative of the US Department of Agriculture’s National Institute of Food and Agriculture under Grant 2019-67013-29297; and the Ohio Agricultural Research and Development Center’s North Central Soybean Research Program under Grants OHO01412 and OHO01355-MRF.

## Data Availability Statement

The data that support the findings of this study are openly available in Ag Data Commons at https://doi.org/10.15482/USDA.ADC/1528082.

## Footnotes

[1] Audacity® software is copyright © 1999-2022 Audacity Team. The name Audacity® is a registered trademark.

