## Supplementary figures and images for "Measuring factors affecting honey bee attraction to soybeans using bioacoustics monitoring"

### Supplemental Figure 1

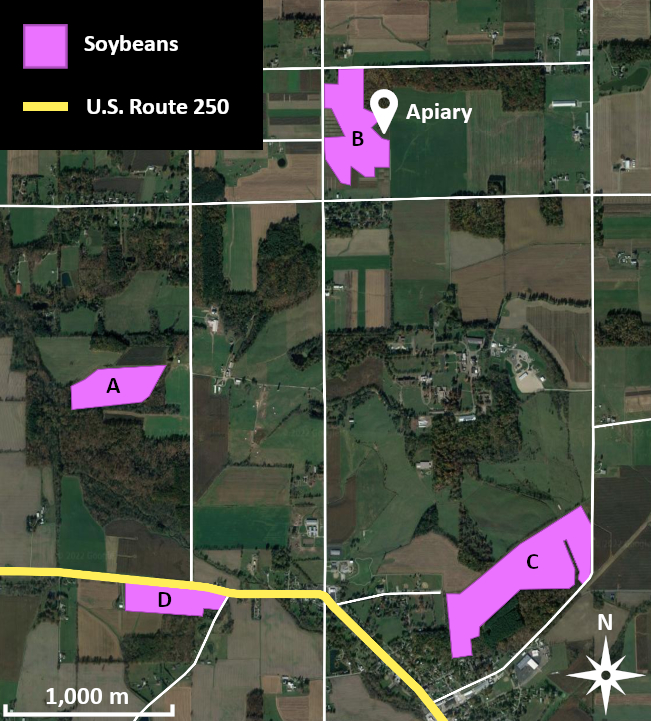

### Supplemental Table 1

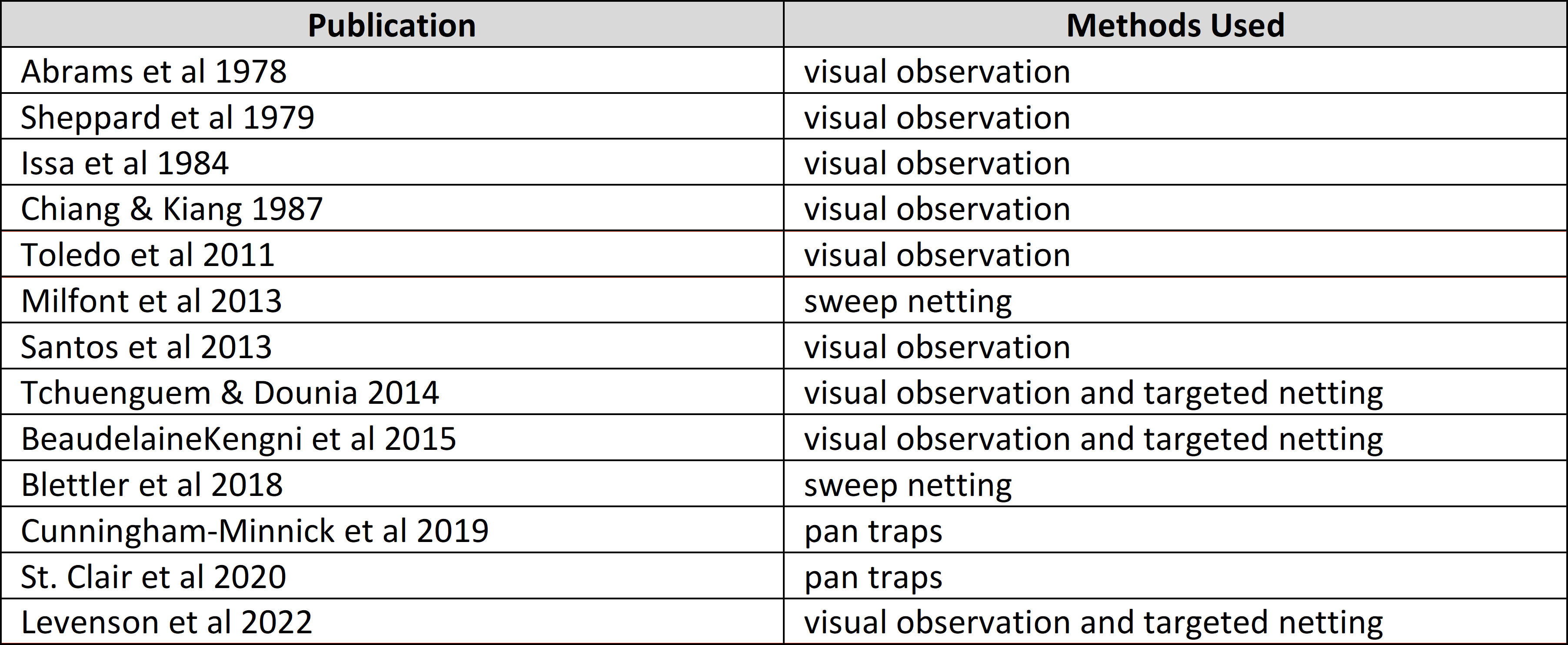

### Supplemental Table 2

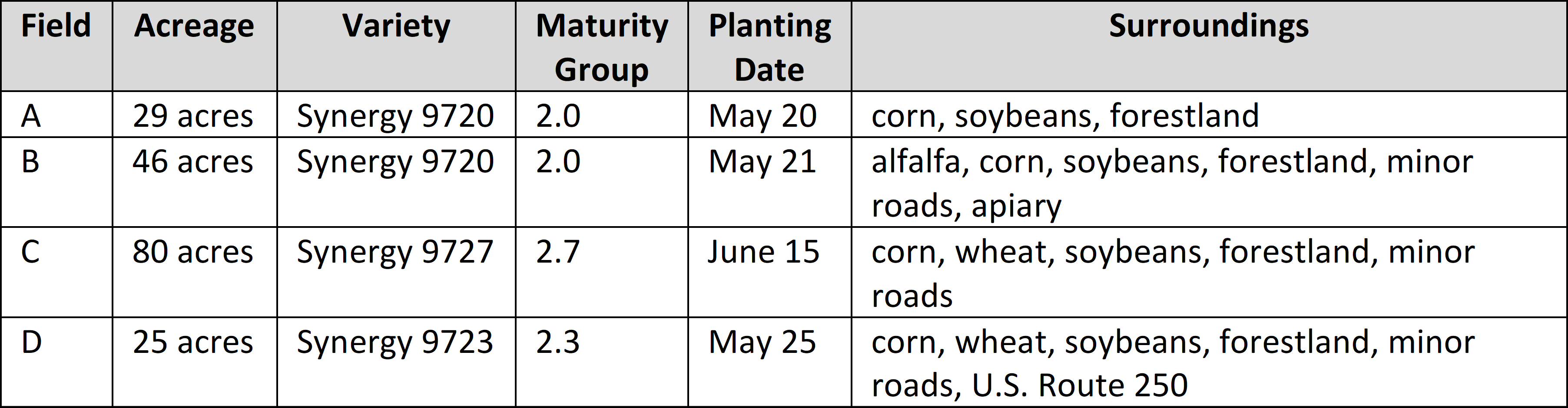

### Supplemental Table 3

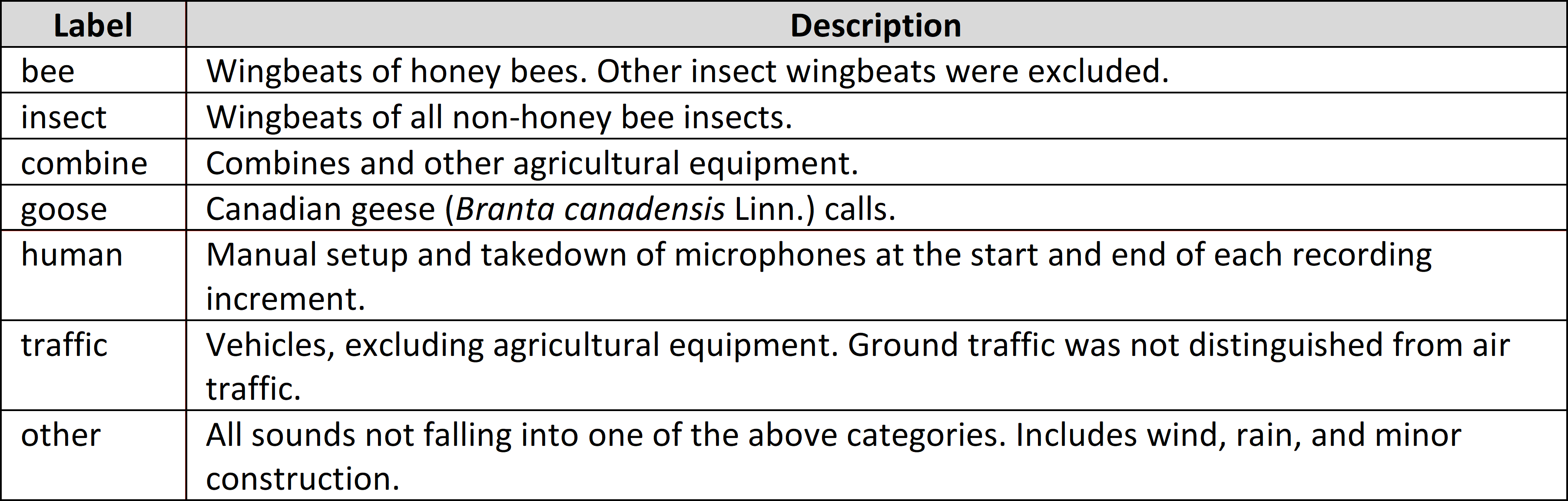

### Supplemental Table 4

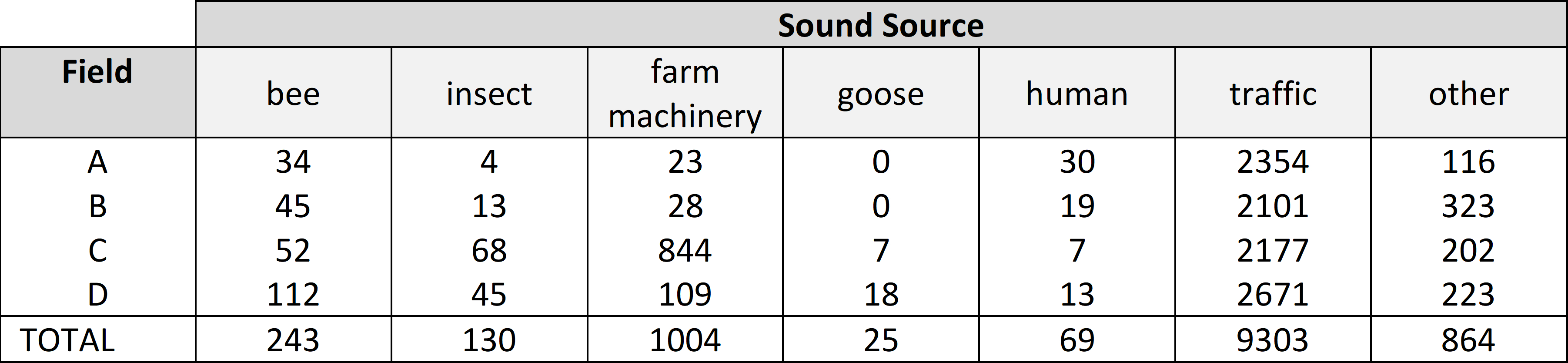
